# Impact of G-tract RNAs and the DHX36 helicase on stress granule composition and formation

**DOI:** 10.1101/2025.06.16.659950

**Authors:** Li Yi Cheng, Nina Ripin, Thomas R. Cech, Roy Parker

## Abstract

Stress granules are RNA-protein condensates that form in response to an increase in untranslating mRNPs. Stress granules form by the condensation of mRNPs through a combination of protein-protein, protein-RNA, and RNA-RNA interactions. Several reports have suggested that G-rich RNA sequences capable of forming G-quadruplexes promote stress granule formation. Here, we provide three observations arguing that G-tracts capable of forming rG4s do not promote mRNAs partitioning into stress granules in human osteosarcoma cells. First, we observed no difference in the accumulation in stress granules of reporter mRNAs with and without G-tracts in their 3’ UTRs. Second, in U-2 OS cell lines with reduced DHX36 expression, which is thought to unwind G-quadruplexes, the partitioning of endogenous mRNAs was independent of their predicted rG4-forming potential. Third, while mRNAs in stress granules initially appeared to have a higher probability of forming rG4s than bulk mRNAs, this effect disappeared when rG4 motif abundance was standardized by mRNA length. However, we observe that in a G3BP1/2 double knockout cell line, reducing DHX36 expression rescued stress granule-like foci formation. This indicates that DHX36 can limit stress granule formation, potentially by unwinding trans rG4s, or limiting other intermolecular RNA-RNA interactions that promote stress granule formation.

**Key Points:** - G-tract RNAs with quadruplex forming potential in an mRNA do not affect its partitioning into stress granules

- mRNA partitioning to stress granules is dependent on mRNA length rather than rG4-forming potential

- DHX36, a DEAH-box helicase that unwinds RNA G-quadruplexes, limits stress granule formation

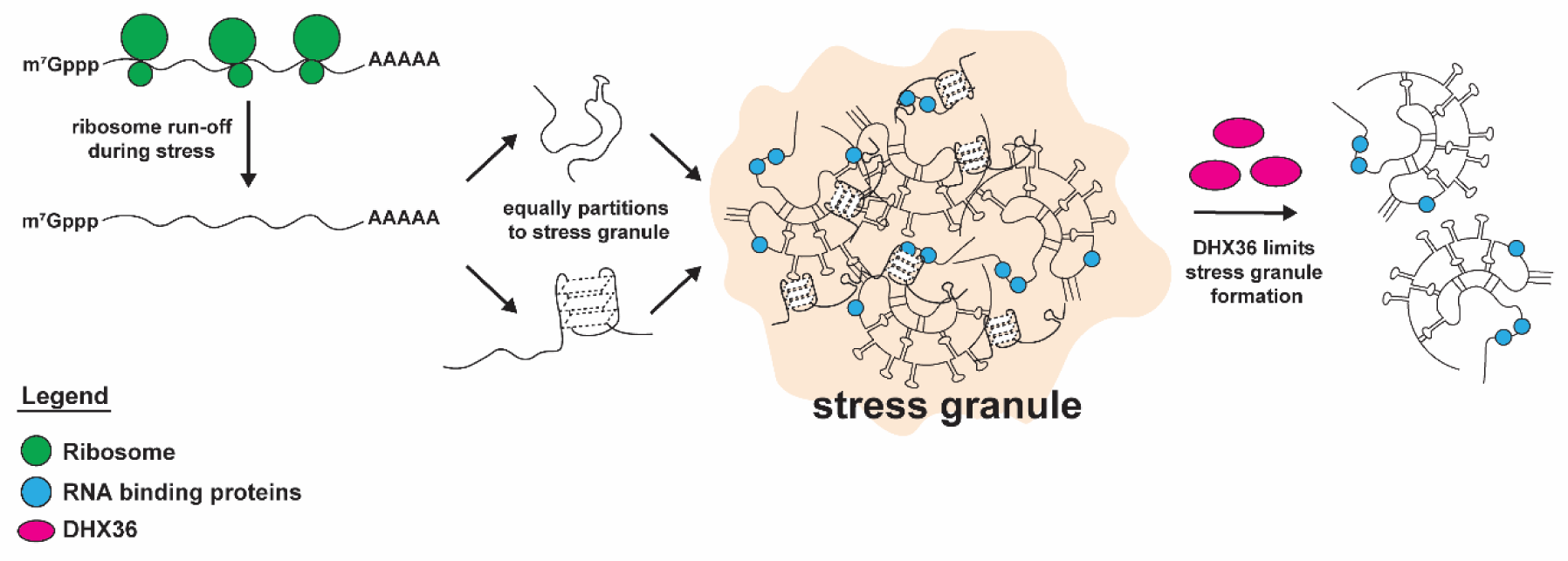

## INTRODUCTION

Stress granules are membrane-less cytoplasmic messenger ribonucleoprotein (mRNP) assemblies that form upon various types of cellular perturbation such as oxidative stress and heat shock. Stress granules are composed of proteins and RNAs, and are thought to form from the summation of protein-protein, protein-RNA and promiscuous RNA-RNA interactions when mRNA translation is inhibited and ribosomes run off mRNAs (1–4). We have previously demonstrated that stress granules contain molecules from essentially all mRNA species found in the cytoplasm, with no single mRNA or ncRNA making up more than 1% of the stress granule RNA content (5). Stress granule formation is promoted by an excess of free untranslating mRNAs (6), and is inhibited by drug treatments that trap ribosomes on mRNAs (7). These observations suggest that concentration driven condensation of ribosome-free and translationally inactive mRNAs form the basis for stress granule assembly.

RNA serves two roles in the formation of stress granules. First, as a critical component of stress granules, RNAs can serve as a scaffold to recruit RNA binding proteins that promote stress granule formation. For example, deletion of the G3BP1 RNA binding domain abolishes stress granule formation in cells (8, 9), suggesting that G3BP1 is required to bind RNA to promote stress granule formation (8–10). RNA binding can also recruit “client proteins”, which are RNA binding proteins that localize to stress granules through binding resident RNAs (11).

In a second role, RNAs can also facilitate stress granule formation through enhancing RNA condensation. *In vitro*, we previously demonstrated that RNA self-assemblies made from total yeast RNA are similar to the stress granule transcriptome (12). Similar work by others have also shown that such RNA self-assemblies can be formed more readily when the trigger RNA species can engage in stable intermolecular interactions (13, 14). Direct evidence for intermolecular RNA-RNA interactions promoting RNP granules comes the observations that specific base-pairing in *trans* between *oskar* mRNAs, or *bicoid* mRNAs, is required for their recruitment to RNP granules during Drosophila oocyte development (15, 16). During cellular stress, mRNA sequences that are normally protected by ribosomes become exposed, increasing their propensity to form promiscuous intermolecular RNA-RNA interactions to enhance mRNP granule formation (17).

One type of RNA structure that might contribute to stress granule formation is the RNA G-quadruplex (rG4).

rG4s are composed of two or more stacked guanine quartets, each formed by the planar interaction of guanine bases via Hoogsteen base pairing. *In vitro*, rG4 sequences can undergo phase separation and form droplets (18, 19). These rG4 assemblies could help to bring together a high local concentration of RNAs and/or rG4 binding proteins to favour assembly formation.

Several studies have proposed that rG4s promote stress granule formation. This was first suggested by the observation that oligonucleotides that can form rG4s can promote RNP condensation *in vitro* and stress granule formation when transiently transfected into cells (13). In addition, several high-throughput transcriptomic studies have revealed numerous putative rG4-forming sequences in mRNAs (20–22). Moreover, mRNAs with putative rG4 sequences were suggested to accumulate in stress granules to a greater extent than mRNAs without predicted rG4 sequences (22). More recently, intermolecular rG4s have been proposed to bind and nucleate proteins that enhance stress granule formation. Specifically, interaction of rG4s with RNA binding proteins such as DNAPTP6 (23) and SERF2 (24) were suggested to coordinate stress granule assembly in mammalian cells.

The possible role of rG4s in stress granules raises the possibility that the DEAH-box helicase DHX36, which is the predominant protein known to resolve rG4s in HeLa cells (25), regulates stress granule formation by targeting and unwinding endogenous rG4s (26). *In vitro* biochemical studies showed that DHX36 exhibits tight binding affinity with rG4s in the sub-nanomolar range (27), and preferentially unwinds rG4s relative to other nucleic acid secondary structures (25, 28). In cells, loss of DHX36 was shown to increase the abundance of G-rich, translationally inactive DHX36 target mRNAs capable of forming rG4s (22). Moreover, DHX36 is found within stress granules upon cellular stresses (22, 29, 30). Precedent for how DHX36 might affect stress granules comes from studies on eIF4A, which can limit stress granule formation by limiting intermolecular RNA-RNA interactions (31). DHX36 might affect stress granule formation by preventing rG4s from forming in mRNAs that serve as binding sites for rG4 binding proteins that promote stress granules. Alternatively, DHX36 could limit the formation of rG4s in trans between multiple RNAs, which then could contribute to intermolecular RNA-RNA interactions that stabilize stress granule assembly.

Herein, we examined if G-rich sequences with rG4-forming potential have any role in targeting mRNAs into stress granules in human U-2 OS cells. We report several key observations. First, mRNA partitioning to stress granules is independent of G-tracts in several tested mRNAs. Second, partitioning of endogenous mRNAs with putative rG4 motifs is unchanged upon DHX36 knockdown, and their localization to stress granules is more length-dependent than rG4-dependent. Third, we observe that DHX36 can limit stress granule formation, perhaps by resolving *trans* rG4 or other types of intermolecular RNA-RNA interactions.

## MATERIALS AND METHODS

### Circular dichroism

RNA was purchased from IDT Technologies and dissolved in double-distilled water. For CD, RNA was then adjusted to 10 mM Tris (pH 7.5) with or without 100 mM KCl or LiCl as indicated. The 150 μL sample (∼ 0.5 mg/mL RNA) was heated to 95°C for 5 min and snap cooled. Either KCl buffer or LiCl buffer alone was used as a baseline control for each salt condition. CD spectra were collected with Chirascan Plus Circular Dichroism and Fluorescence Spectrometer (Applied Photophysics) using a 0.5 mm path length cell. Spectra were recorded from 360 to 200 nm with a 0.5 nm step size and 0.5 sec integration time. When switching between samples, the cuvette was washed 2× with H2O and once with 100% ethanol. Baselines were subtracted from experiment spectra and circular dichroism was converted to mean residue molar ellipticity. Spectra are reported in mean residue molar ellipticity.

### Cell culture

Human osteosarcoma U-2 OS cells were maintained in Dulbecco’s modified Eagle’s medium supplemented with 10% fetal bovine serum and 1% penicillin/streptomycin at 37°C/5% CO_2_.

### Cell lines and plasmids

Wild-type and G3BP1/2 double knock-out (dKO) U-2 OS are kindly provided by Paul Anderson’s Lab at Brigham and Women’s Hospital, Boston, MA, USA (8). Tet-inducible luciferase reporter was a gift from Moritoshi Sato (Addgene plasmid #64127; http://n2t.net/ addgene:64127; RRID:Addgene_64127, (32).

Various rG4 sequences were synthesized by GenScript (Table S1 for sequences) and cloned into the 3’ UTR of the luciferase reporter via InFusion cloning. 1x reporter constructs are dual luciferase reporter constructs with FLuc as a control. Both RLuc and FLuc are controlled by the Tet-inducible promoter, independently. 5x reporter constructs only has RLuc at the 5’ of rG4 motifs, it does not contain FLuc as an internal control.

To integrate the luciferase reporter constructs into the AAVS1 safe harbour locus, WT U-2 OS cells were transfected with 1 μg CRISPR/Cas9 plasmid (pRP2854) in conjunction with 1 μg appropriate luciferase reporter construct using JetPRIME transfection reagent (VWR Scientific, 89129-922). Transfection of pRP2854 alone was used as a negative control. Twenty-four hours following transfection, cells were split from a six-well plate to a T25 flask with media containing 1 μg/mL puromycin (Sigma-Aldrich, P9620) to begin selection for cells with genomic integration. Following 24 h with puromycin selection, media was replaced with fresh puromycin. After all cells were dead in the negative control plate, media was replaced with fresh media lacking puromycin for 48 h. This is an optional step to help get rid of any residual plasmid. Puromycin was then added for another 48 h to finalize the selection. Single colony selection was not done for these experiments.

DHX36 hypomorph (DHX36hm) constructs in WT and G3BP1/d dKO U-2 OS cell lines were generated using methods from (33). Briefly, two CRISPR/Cas9 guide RNAs targeting different regions within the DHX36 locus were designed using the Integrated DNA Technologies (IDT) CRISPR guide target design tool. Overlapping oligos (DHX36 sgRNA 1, 2 sense and DHX36 sgRNA 1,2 antisense [Table S1]) were annealed in T4 DNA ligase buffer (NEB, B0202S) and ligated into the BbsI-HF (NEB, R3539S) sites in pSpCas9(BB)-2A-GFP (px458) (48138; Addgene) using T4 DNA ligase (NEB, B0202S). To generate DHX36hm in U-2 OS and G3BP1/2 KO U-2 OS lines, cells (T-25 flask; 60% confluent) were co-transfected with 3 µg pSpCas9(BB)-2A-GFP-DHX36 sgRNA1+2 and 400 ng of pcDNA3.1-puro using 15 µl of Lipofectamine 2000 (Thermo Fisher Scientific, 11668019) according to the manufacturer’s instructions. Twenty-four hours following transfection, cells were puromycin selected as described above. Selective medium was replaced with normal growth medium after all cells were dead in the negative control. When cells became 80% confluent, cells were serial diluted and plated on 15 cm dishes. After visible colony formation, individual colonies were isolated, propagated, and tested for DHX36 loss via Immunoblotting.

Generation of the pLenti-EF1-Blast-HA-FLAG-DHX36 lentiviral plasmids was performed as described in (34). Briefly, untagged DHX36 sequences were amplified from Addgene (Addgene plasmid #159585, #159587), and the sequences were inserted into the XhoI/XbaI sites of pLenti–EF1-Blast vector using In-Fusion seamless cloning (Takara Bio).

Subsequently, synonymous mutations, designed via the Synonymous Mutation Generator (35), were introduced into HA-FLAG-tagged DHX36 domain constructs to make siDHX36-resistant sequences using PCR and InFusion cloning (Takara Bio). 3 out of 4 sites were made resistant to the pooled siDHX36 purchased from Dharmacon. Subsequently, siRNA-resistant DHX36 domain constructs were generated using a lentivirus system. HEK293T cells (T-25 flask; 80% confluent) were co-transfected with 2.7 µg pLenti-EF1-Blast-HA-FLAG-DHX36, 870 ng of pVSV-G, 725 ng of pRSV-Rev, and 1.4 µg of pMDLg-pRRE (36), using 20 µl of Lipofectamine 2000. The medium was collected 48 h after transfection and filter-sterilized with a 0.45 µm filter. Then, U-2 OS G3BP1/2dKO + DHX36hm cells (T-25 flask; 80% confluent) were transduced with 1 ml of lentiviral stocks containing 10 µg/ml of polybrene (Millipore Sigma, TR-1003-G) for 1 h. DMEM was then added to the flask after an hour. 24 h after transduction, cells were reseeded into a T-75 flask containing 10 µg/ml blasticidin (Thermo Fisher Scientific, A11139-03) selective medium. Cells were maintained in selective medium for 4–5 days before returning to normal DMEM.

### Immunoblotting

Cells were washed with ice-cold phosphate-buffered saline (PBS) and lysed with Pierce RIPA Buffer (Thermo Fisher Scientific, 89900), 1* Phosstop™-phosphatase inhibitor (Roche, 4906837001), and 1* cOmplete Mini EDTA-free Protease Inhibitor Cocktail (Sigma-Aldrich, 11836170001). Cells were lysed on ice for 15 min (flicked every 5 min) and then clarified by centrifugation at 4°C, max speed for 10 min. 4× Nu-PAGE LDS sample buffer (Thermo Fisher Scientific, NP0007) was added to lysates to a final concentration of 1×, samples were boiled for 10 min at 70°C, and then loaded into 4–12% Bis-Tris Nu-PAGE gel (Thermo Fisher Scientific, NP0336BOX) and transferred with Iblot™2 PVDF transfer stacks (Thermo Fisher Scientific, IB24002). Membranes were blocked with 5% nonfat-dried milk in Tris-buffered saline with 0.1% Tween-20 (Sigma-Aldrich, P9416) (TBST) for 1 h and then incubated with primary antibody in TBST for 1 h at room temperature or overnight at 4°C. Antibody dilutions are listed in Table S1. Membranes were washed 3× with TBST and then incubated with secondary antibody at room temperature for 1 h in TBST. Membranes were washed 3× again in TBST, and antibody detection was achieved with chemiluminescence substrate (Thermo Fisher Scientific, 34095).

### Stress conditions

To induce stress granules, cells were treated with 500 µM sodium arsenite (Sigma-Aldrich S7400) for 1 h at 37°C unless otherwise specified. Cells were fixed after the completion of stress with 4% paraformaldehyde. To examine stress recovery, cells were incubated with 300 µM sodium arsenite for 1 h before replacing with normal DMEM.

### Dual luciferase assay

U-2 OS cells were plated at 50% confluency in 96-well plates for luciferase assay. On the next day, cells were lysed using the Passive Lysis Buffer (Promega). Renilla and firefly luciferase activities were measured using the Dual-luciferase Reporter Assay (Promega) as per manufacturer’s protocol with the CLARIOstar® Plus Multi-mode Microplate Reader. Wild-type U-2 OS cells with no reporter constructs were used as the negative control for background luciferase activity. In all cases, renilla luciferase values were normalized to firefly luciferase values.

### siRNA-mediated knockdown

For siRNA knockdowns, 200,000 cells per well were seeded into six-well plates. After 24 h, cells were transfected with 20 nM siGENOME SMARTpool siDHX36 (Dharmacon, M-013167-00-0005) using Lipofectamine RNAiMAX (Invitrogen, 13778-150). For each reaction, 5 µl Lipofectamine was added to 150 µl Opti-MEM Medium. 20 nM siRNA was added to another tube with 150 µl Opti-MEM Medium. Both were combined, vortexed, and incubated for 20 min at RT. 300 µl siRNA-Lipofectamine mix was added per well with 1.7 ml media. 24 h after transfection, cells were trypsinized and seeded onto a glass coverslip in 24-well plates to be fixed for imaging or onto 6-well plates to be lysed for Western blots the day after.

### smFISH probes

Custom single-molecule fluorescence in situ hybridization (smFISH) probes against WAC, NAA50 (ENST00000240922.7), PURB (ENST00000395699.3) and SLMO2 (ENST00000355937.8) were designed with Stellaris RNA FISH Probe Designer, and labeled with Quasar 670 dye based on a protocol described in (37). Oligo d(T)30-Cy3 probes were purchased from IDT.

smiFISH probes for RLuc were generated as described in (38) with minor adjustments. RLuc smiFISH probes were designed with the Biosearch Technologies Stellaris Probe Designer version, 4.2. Both probes and the secondary FLAPY sequence were ordered from IDT. Primary probes were pooled together in equimolar amounts to a final concentration of 100 µM total oligo.

### Sequential IF and FISH

Sequential immunofluorescence and smFISH/smiFISH on fixed U-2 OS cells were performed with homemade buffers (39) according to the manufacturer’s protocol: (https://biosearchassets.blob.core.windows.net/assets/bti_custom_stellaris_immunofluorescence_seq_protocol.pdf). Briefly, U-2 OS cells were seeded on sterilized coverslips in 24-well tissue culture plates. At ∼80% confluency, media was exchanged 1 h before experimentation with fresh media. After stressing cells (see section “Stress conditions”), the media was aspirated and the cells were washed with 1× PBS pre-warmed to 37°C. The cells were fixed with 350 μL 4% paraformaldehyde for 11 min at room temperature. After fixation, cells were washed thrice with 1× PBS, permeabilized in 0.1% Triton X-100 in 1× PBS for 5 min and washed once with 1× PBS. For IF detection, coverslips were incubated in primary antibody for 1 h at room temperature. Coverslips were washed three times with 1× PBS. Then cells were incubated in secondary antibody for 1 h at room temperature. Again, coverslips were washed three times with 1× PBS. Subsequently, cells were fixed again with 350 μL 4% paraformaldehyde for 11 min at room temperature and washed thrice with 1x PBS. Cells were then treated with smFISH Buffer A for 5 min. Coverslips were transferred to a humidifying chamber with smFISH probes and placed in the dark at 37°C for 16 h. Coverslips were placed in Buffer A for 30 min in the dark, replaced with fresh Buffer A for another 30 min in the dark, washed twice with Buffer B for 5 min each and placed onto a slide with VECTASHIELD Antifade Mounting Medium with DAPI (Vector Labs, H-1200).

In order to maintain consistency, the same protocol was utilized in IF only experiments; however, the portions of the protocol calling for smFISH were omitted. Coverslips were mounted with Prolong Glass Antifade Mountant with NucBlue Stain (Thermo Fisher Scientific, P36981). Antibodies used are listed in Table S1.

### Microscopy

Majority of fixed U-2 OS cells stained by immunofluorescence and smFISH were imaged using the inverted Nikon Ti2 Eclipse spinning disk confocal microscope with either a 60x NA 1.27 water immersion objective or a 100x NA 1.45 oil immersion objective and a Hamamatsu ORCA Fusion BT sCMOS camera. Immunofluorescence of smFISH for NAA50, PURB and SLMO2 were imaged using the inverted Nikon Ti Eclipse spinning disk confocal microscope with a 100× NA 1.4 oil immersion objective and a 2× Andor Ultra 888 EMCCD camera (BioFrontiers Advanced Light Microscopy Core). More than three images were recorded at room temperature per biological replicate, with z-sections were taken for each experiment. Every experiment was performed in two or three biological replicates.

### Image analysis and quantification

Image processing was conducted using Fiji image processing package, with all shown images being the maximum intensity projection of a series of z-sections. Image segmentation masks, quantification of fluorescence intensities, SG counts, smFISH spots and cell measurements were obtained using CellProfiler 4.2.5. Partition coefficients were determined by the number of FISHspots in granule/cytoplasm, and analyzed using R-studio. To count cells with stress granule like foci, images were blinded and manually examined.

### Statistical analysis

Unpaired, two-sided *t* test was performed on the mean of all biological replicates. In experiments with more than two groups, one-way ANOVA with Tukey’s multiple comparisons test was performed and adjusted *p*-value reported, unless stated otherwise. All statistical analysis was performed in Graphpad Prism 8.0.

## RESULTS

### Generation of a luciferase reporter system to probe stress granule partitioning

rG4s in mRNAs have been proposed to promote stress granule partitioning due to their phase separation properties *in vitro* (12, 13). To test this possibility, we created reporter mRNAs with and without G-tracts inserted into the 3’ UTR and then examined if putative rG4 motifs affected the localization of the reporter into stress granules. We constructed a set of luciferase reporters with either predicted rG4 forming sequences or non-rG4 forming sequences of different lengths (1x or 5x) in their 3’ UTRs (Fig. 1A, Table S1). Reporter constructs included a Tetracycline-on expression system so that the RNAs should only be expressed in the presence of doxycycline (abbreviated as Dox in figures). These constructs were targeted to the AAVS1 safe harbour locus for integration and stable expression following transfection in wild-type (WT) U-2 OS cells using CRISPR/Cas9 genome editing and a donor template DNA. However, it should be noted that whether our reporter constructs form stable rG4 structures in cells remains uncertain, as cellular conditions may favour unfolding by RNA helicases (40, 41) and/or promote the formation of alternative, competing secondary structures.

**Figure 1.**
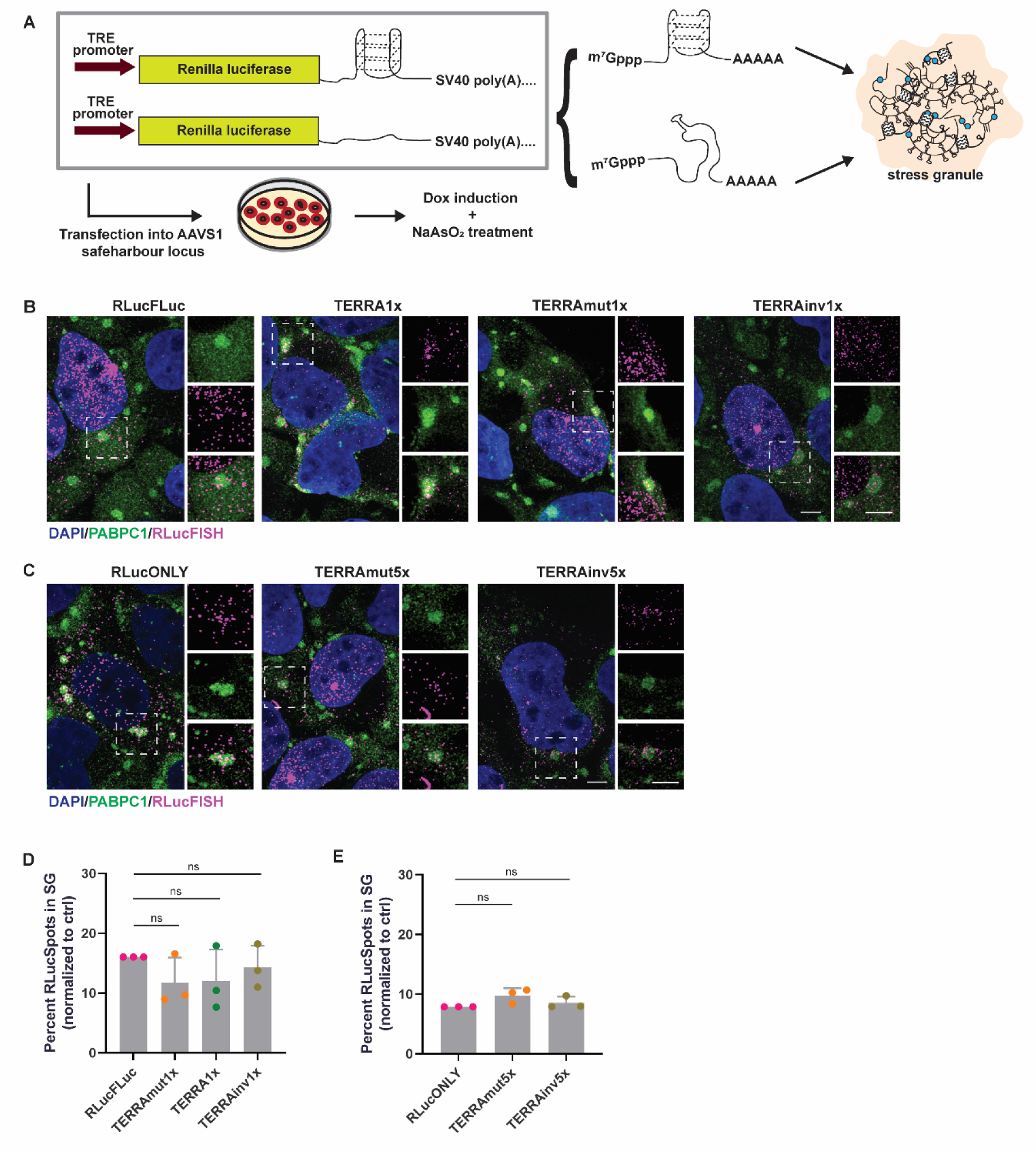
3’ UTR G-tracts do not alter mRNA localization into stress granules. (A) Schematic of rG4 reporter localization. To determine if mRNA with a rG4 motif will be preferentially recruited into stress granules compared to a mRNA without a rG4 motif, sequences either predicted to form rG4s or controls defective in rG4 formation are inserted into the 3’ UTR of Renilla luciferase (Rluc). The reporter is under the control of a Tetracycline-inducible system where addition of doxycycline will activate the TRE promoter and transcribe the downstream reporter mRNA. Reporter constructs are stably transfected into the AAVS1 locus of U-2 OS cells. Blue circle: RNA binding protein. (B-C) Immunofluorescence of DAPI (blue), PABPC1 (green), and RLuc RNA (magenta) in U-2 OS cells stably expressing (B) 1x reporter constructs and (C) 5x reporter constructs treated with 500 µM NaAsO_2_ for 60 min. Scale bar = 5 µm. (D-E) Quantification of RLuc RNA FISH spots in stress granules as in (B) and (C) respectively, normalized to no 3’ UTR control. Data analyzed with one-way ANOVA, corrected with Tukey’s multiple comparisons test and represented as mean ± s.d.. ns = non-significant, p > 0.05. Three biological replicates quantified. TERRA reporter sequences in Table S1.

We introduced a specific set of sequence motifs, computationally predicted to adopt rG4 conformations, into the 3’ UTR. One of the best understood ncRNAs that forms rG4s is the telomeric repeat-containing RNA (TERRA) (42–44). However, since TERRA is usually localized in the nucleus and might be involved in telomere-related function, we generated a construct containing an inverted TERRA sequence (Table S1), which would also be predicted to form a rG4 (referred to as TERRAinv). Using circular dichroism, we confirmed that the 24-mer TERRAinv formed similar rG4 structures to that of TERRA in the presence of K+ *in vitro* with a distinguishing positive peak at 210 nm (Supplementary Fig. 1A) while rG4 signatures were abolished in Li+ solution (Supplementary Fig. 1B). As a control, we generated a construct that contains the same G-content as the rG4 motifs but lacks consecutive guanine bases. This construct, referred to as TERRAmut, has reduced rG4-forming capability, as confirmed via circular dichroism where it showed a negative peak at at 210 nm that is indicative of A-form RNA rather than rG4 (45) (Supplementary Fig. 1A & 1B).

We validated that all reporter constructs were successfully integrated into the AAVS1 locus using primers flanking Renilla luciferase (RLuc) (Supplementary Fig. 1C shown for 1x constructs, data for 5x constructs not shown). Next, we showed in RLucFLuc U-2 OS cells that RLuc and Firefly luciferase proteins were expressed only upon doxycycline induction and were enzymatically active, demonstrating the mRNAs were fully functional (Supplementary Fig. 1D & 1E).

Together, this shows that our luciferase reporter system was functional in cells and allowed the expression of functional mRNAs.

### 3’ UTR G-tracts do not alter mRNA localization into stress granules

To determine whether G-tracts in 3’ UTRs affect mRNA localization in stress granules, we examined the localization of our reporter mRNAs using immunofluorescence. We induced expression of the reporter constructs containing G-tracts with the potential of forming a single rG4 motif by treating cells with doxycycline for 48 hours, followed by exposure to NaAsO_2_-induced oxidative stress for one hour. We then imaged the localization of RLuc reporter mRNAs in the cell with single-molecule inexpensive fluorescence in situ hybridization (smiFISH), which uses unlabelled gene-specific probes with a FLAP sequence that is subsequently recognised by the fluorescently labelled secondary detector probes (38, 46). To confirm that the smiFISH signal was real, we also stained cells for their FLuc and RLuc protein expression. We showed that cells with RLucFISH signals are also positive for either FLuc (Fig. 1B & Supplementary Fig. 2A) or RLuc (Fig. 1C & Supplementary Fig. 2B).

The fraction of the reporter mRNAs localized to stress granules was then determined by quantification of the number of smFISH spots overlapping with PABPC1+ stress granules, compared to the total number of cytoplasmic mRNAs (see Methods).

We observed no statistical difference in the fraction of reporter mRNAs accumulating in stress granules with or without the insertion of 3’ UTR G-tracts. Specifically, we observed 11.8 ± 4.3% and 14.2 ± 2.8% (throughout this paper, errors are S.D. and the number of biological replicates is given in the figure legends) of stress granules contained TERRA and TERRAinv reporter mRNAs, respectively, while we saw 11.5 ± 3.2% of stress granules contained TERRAmut mRNA (Fig 1B & 1D). Thus, these experiments did not support the preferential enrichment of rG4 sequence motifs in stress granules.

In addition to our artificial 3’ UTR TERRA sequences, we also inserted the 3’ UTR rG4 sequence motifs from the endogenous mRNA, *APP*, into our reporter system. The *APP* 3’ UTR rG4 sequence has been previously validated to form rG4 structures (47). Similar to the TERRA constructs, we did not observe differences in reporter mRNA partitioning to stress granules in the presence of 3’ UTR APP rG4 sequence motifs (Supplementary Fig. 2C & 2D). Thus, having 3’ UTR G-tracts that are predicted to form a single rG4 in an mRNA did not alter its recruitment into stress granules.

The assembly of macromolecules into condensates is generally proportional to the number of interaction sites (11). Indeed, we have previously demonstrated that mRNA partitioning into stress granules can be enhanced by increasing protein binding sites on the mRNA in a dose dependent manner (48). Given this, we considered the possibility that the contribution of a single rG4 sequence motif in an mRNA to stress granule partitioning might be very small and difficult to detect. We therefore tested reporter constructs harbouring five repeats of the putative rG4s in the 3’ UTR to determine if adding more rG4 sequence motifs could alter mRNA localization to stress granules.

We observed that having five putative rG4-forming sequence motifs at the 3’ UTR also did not increase reporter mRNA partitioning to stress granules (Fig. 1C & 1E). Taken together, this suggests that the presence of one or more 3’ UTR rG4 sequence motifs does not affect mRNA localization into stress granules in U-2 OS cells.

### rG4 sequence motifs within endogenous mRNAs do not predict stress granule partitioning

DHX36 is a member of the DEAH-box helicase family that has been proposed to tightly bind rG4s and unwind them (25, 27, 49). Given this role, we considered the possibility that DHX36 is unwinding rG4s in cytoplasmic mRNAs and thereby preventing rG4s from promoting mRNA accumulation in stress granule. To test this model, we first investigated DHX36 localization in U-2 OS WT cells using immunofluorescence. Consistent with previous results, DHX36 is mostly cytoplasmic, and upon arsenite stress, DHX36 is enriched in stress granules as indicated by colocalization with PABPC1, and therefore could affect RNA structures within stress granules (Supplementary Fig. 3C) (22, 29, 30).

We examined if DHX36 affected the partitioning of endogenous mRNAs with rG4-forming potential. Previous work suggested that DHX36 mRNA targets were enriched in stress granules (22). However, length is also a key determinant for mRNA enrichment to stress granules (5) due to increased multivalency (48). Here, we cross-referenced the DHX36-E335A PAR-CLIP data (22) with the stress granule transcriptome (5) and binned DHX36 target mRNAs in accordance with the number of cross-linked reads per target mRNA normalized by overall mRNA abundance (normalized crosslinked reads per million, NXPM) obtained by DHX36-E335A PAR-CLIP (22).

Our analyses suggest that the enrichment of DHX36 targets in stress granules is likely a result of longer mRNAs having an enrichment of DHX36 targets. Notably, we observed that while RNAs with increasing DHX36 binding sites are more likely to be enriched in stress granules (Fig. 2A, left), these RNAs also tend to be longer in length (Fig. 2A, right). A similar result was observed when integrating data from transcripts harbouring rG4s (20) (Fig. 2B). In this second dataset, the presence of rG4s was defined by a reverse transcriptase stalling (RTS) value > 0.2 (20) which means that > 20% of the reads containing the rG4 motif showed stalling. Thus, a value of 0.2 suggests that the rG4 structure was present in 20% of the population of a particular transcript. The comparison of transcripts without rG4s and transcripts with >20% rG4 shows that longer transcripts show increased RTS values (Fig. 2B, right), which again correlates with stress granule enrichment (Fig. 2B, left). Thus, our analyses are consistent with mRNA length, rather than the presence of rG4 motif in a mRNA, dictating a mRNA’s enrichment to stress granules.

**Figure 2.**
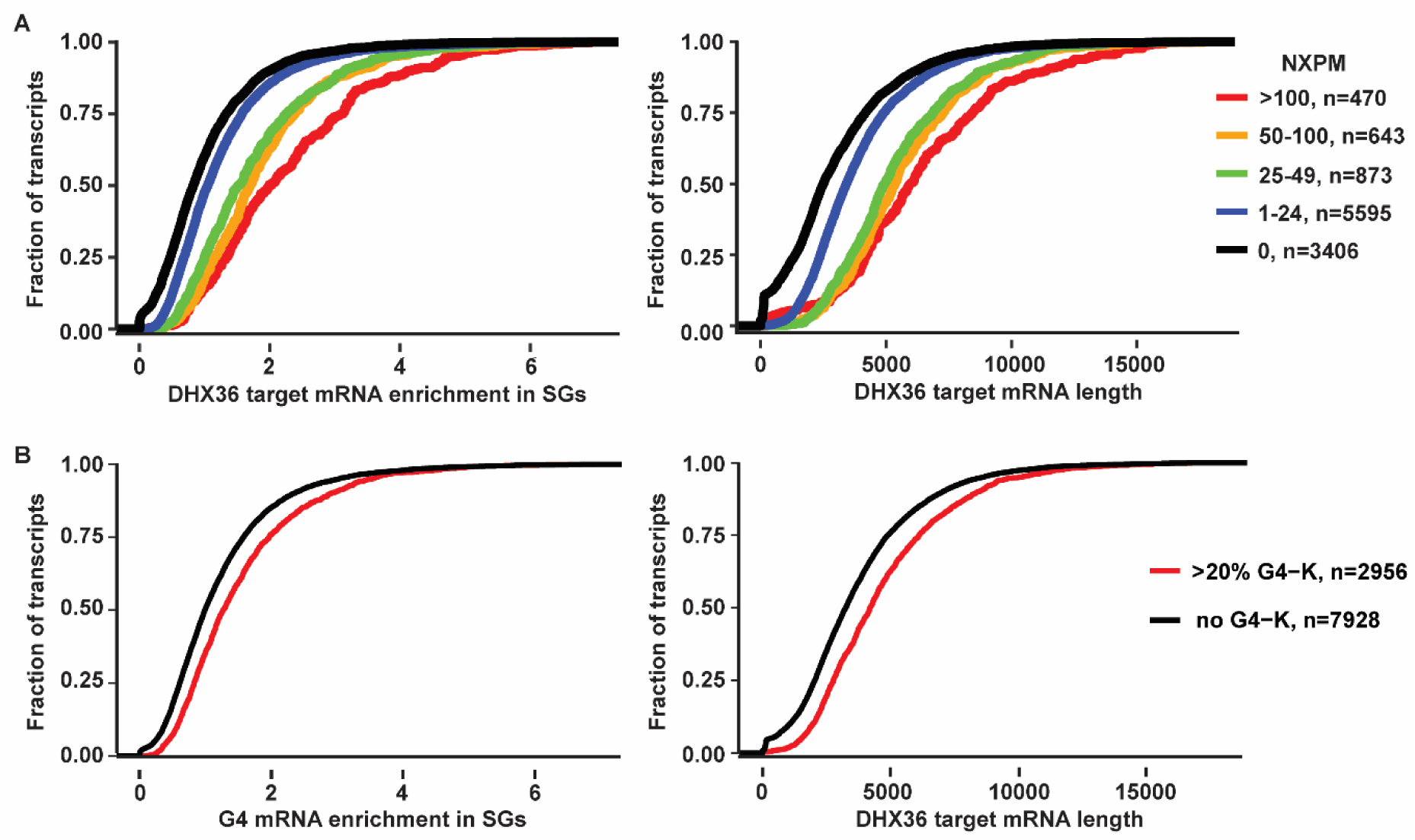
Enrichment of rG4 motifs in stress granule-localized mRNA appears to be a consequence of mRNA length. (A) Cumulative distribution functions showing (left) enrichment in stress granules of DHX36 target mRNAs identified in DHX36-E335A PAR-CLIP (22) compared to non-target mRNAs and (right) length distribution of DHX36 target mRNAs compared to non-target mRNAs. DHX36 target mRNAs are binned in accordance to the number of normalized crosslinked reads per million (NXPM), as described in (22). (B) Cumulative distribution functions showing (left) enrichment of mRNA transcripts harbouring G4 motifs in K^+^ identified previously in (20) compared to non rG4-harbouring transcripts and (right) length distribution of G4-harbouring mRNA transcripts compared to non-rG4 harbouring transcripts. Data on mRNA enrichment in stress granules is obtained from (5).

We then experimentally examined if the loss of DHX36 increased the partitioning of DHX36 target mRNAs with rG4 motifs into stress granules. We used Cas9 targeting the DHX36 gene with two guide RNAs to obtain cell lines with reduced DHX36 protein levels. We obtained colonies with stably reduced DHX36 expression in WT cells (hereafter DHX36hm (for hypomorph)) as shown by western blotting (Supplementary Fig. 3A, 3B left). No colonies were found with a complete loss of DHX36, arguing that DHX36 is essential in U-2 OS cells. As reported in HEK293 cells (22), DHX36hm cells displayed slower growth rate and different morphology due to inefficient spreading in the culture dish (not shown).

Since DHX36 KO has been reported to produce spontaneous stress granules by activating protein kinase R (PKR) in the integrated stress response pathway (22), we examined stress granule formation and PKR phosphorylation in our stable cell lines without stress induction. We did not observe spontaneous stress granules in our DHX36hm cell lines in unstressed conditions, as indicated by PABPC1 IF (Supplementary Fig. 3C). Western blot analyses further corroborated that in the absence of stress, the levels of phospho-PKR and phosphor-eIF2α in DHX36hm did not increase (Supplementary Fig. 3D-F). It should be noted that we might not observe the activation of PKR and spontaneous stress granule formation either because we are working with a hypomorphic DHX36 cell line, better stress recovery in stable cell lines, or because of differences between U-2 OS and HEK293 cells.

Using these stable DHX36hm cell lines, we performed smFISH and quantified the fraction enrichment of DHX36 target mRNAs in stress granules. In this experiment, we designed and tested FISH probes for the DHX36 targets: WAC, NAA50, PURB and SLMO2, that were among the top 100 DHX36 targets in the PAR-CLIP analysis (22). This experiment provided two observations, revealing how DHX36 affects mRNA levels and localization.

First, in U-2 OS DHX36hm cells, we saw the number of mRNA smFISH spots for candidate DHX36 target mRNAs increased (Fig. 3A & Supplementary Fig. 4A), This increase is expected based on the decreased mRNA decay observed for these mRNAs in DHX36 KO HEK293 cells (22).

**Figure 3.**
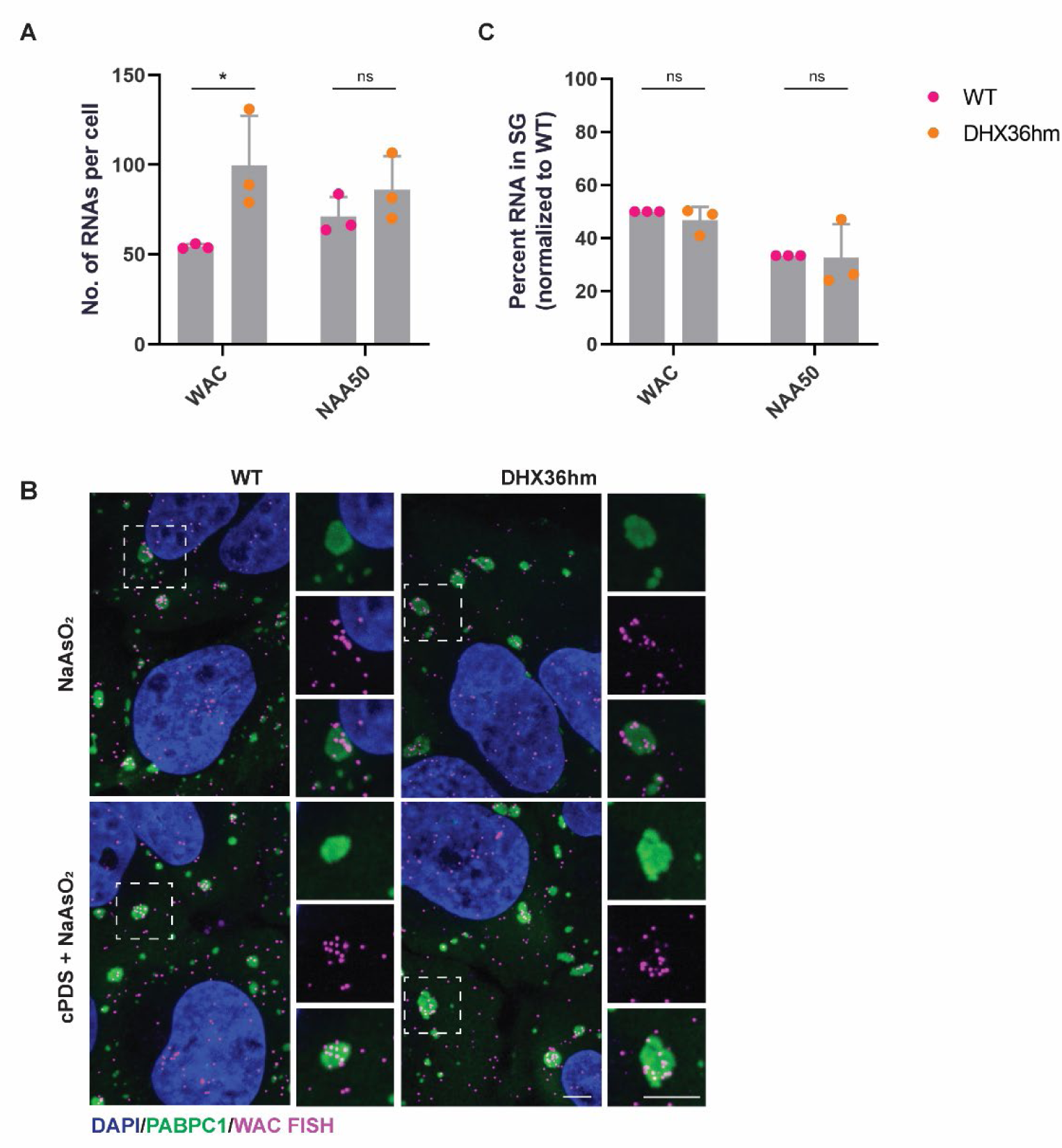
Stress granule localization of mRNAs with rG4 motifs is DHX36-independent. (A) Quantification of WAC and NAA50 RNA FISH spots per cell. Data analyzed with two-tailed unpaired t-test and represented as mean ± s.d. (\**P* = 0.0467). ns = non-significant, p > 0.05. Three biological replicates quantified. (B) Immunofluorescence of DAPI (blue), PABPC1 (green) and WAC RNA (magenta) in WT and stable DHX36hm U-2 OS cells treated with either 500 µM NaAsO_2_ for 60 min, or with 2 µM cPDS for 24 hr and 500 µM NaAsO_2_ for 60 min. Scale bar = 5 µm. (C) Quantification of percent WAC and NAA50 RNA enrichment in stress granules. Data analyzed with two-tailed unpaired t-test, and represented as mean ± s.d.. ns = non-significant, p > 0.05. Three biological replicates quantified.

Second, we observed that the fraction of candidate DHX36 target mRNAs localized to stress granules was the same in WT and DHX36hm cells (Fig. 3B, 3C & Supplementary Fig. 4B & 4C). This argues that DHX36 does not play a major role in modulating the partitioning of these endogenous mRNAs into stress granules.

We also examined the localization of WAC mRNA in the presence of carboxypyridostatin (cPDS), a small molecule that stabilizes rG4s (Fig. 3B). However, upon oxidative stress, the percent enrichment in stress granules remained unchanged (Supplementary Fig. 4E), supporting the hypothesis that the rG4 sequence motifs in mRNAs do not affect mRNA partitioning into stress granules.

Taken together, our results suggest that DHX36 does not affect RNA enrichment into stress granules and that partitioning of endogenous mRNAs with rG4 sequence motifs into stress granules is independent of DHX36.

### DHX36 depletion facilitates stress granule assembly and slows disassembly

Previous work has described eIF4A and DDX6 as DEAD-box RNA helicases that function as “RNA chaperones” to limit stress granule assembly (31, 33). We considered the possibility that DHX36 might function as a part of this RNA chaperone network by altering the protein composition of mRNPs and/or by altering RNA structure.

To understand if DHX36 can affect stress granule dynamics, we first examined changes in stress granule assembly and disassembly kinetics when DHX36 expression was dampened. We assessed stress granule assembly kinetics by exposing cells with low amounts of NaAsO_2_ (100 µM) to allow a sensitive measurement on stress granule formation. We measured the rate of stress granule formation over time using immunofluorescence against PABPC1 and looked at the percentage of cells with stress granules. We observed that stress granules reproducibly formed at a slightly faster rate in DHX36hm cells compared to WT cells (Fig. 4A & 4B). This difference was particularly appreciable at 45 min and 60 min post-NaAsO_2_ treatment (Fig. 4B, 4D & 4E).

**Figure 4.**
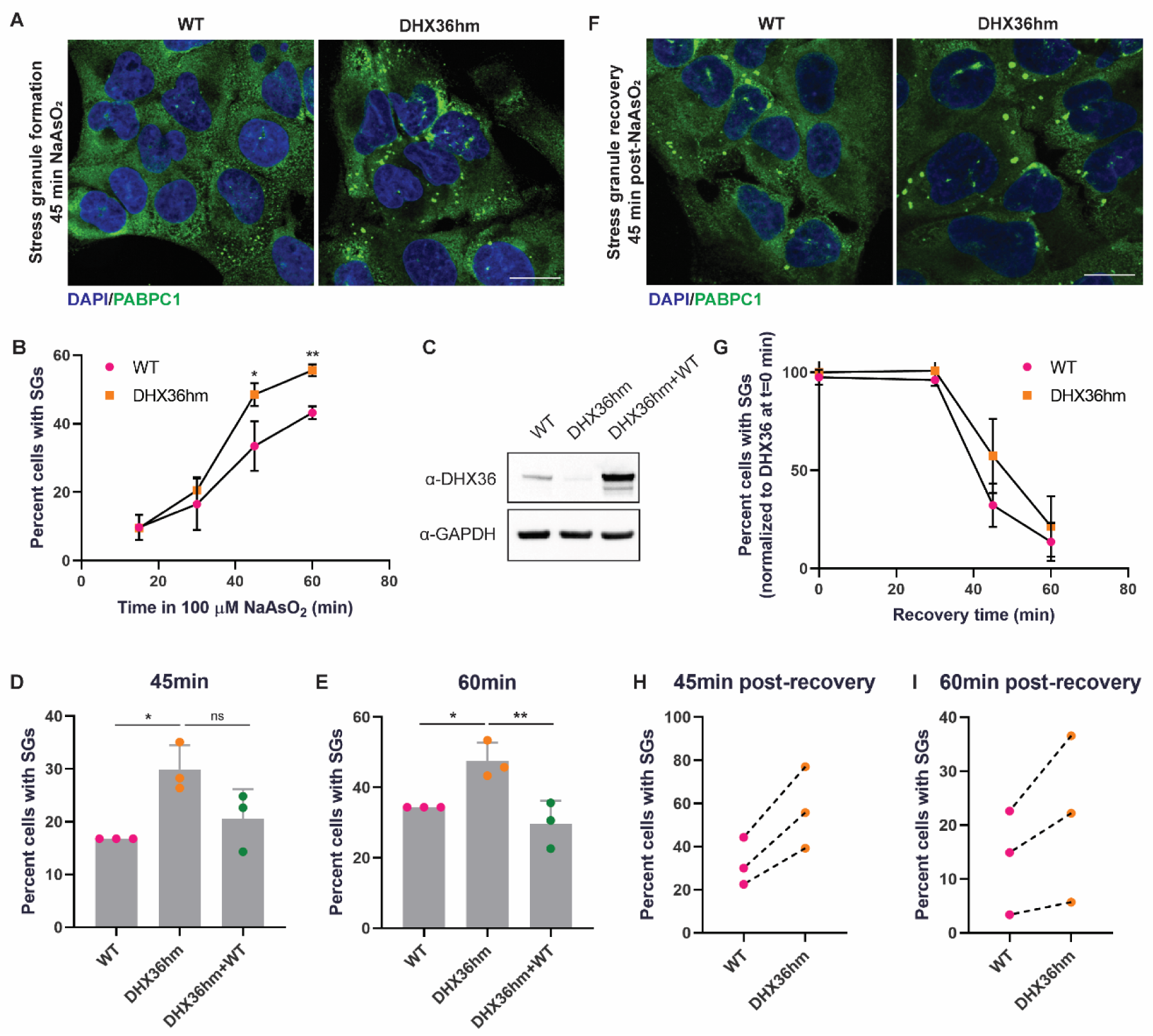
DHX36 depletion facilitates stress granule assembly and slows disassembly. (A) Immunofluorescence of DAPI (blue) and PABPC1 (green) in WT and stable DHX36hm U-2 OS cells treated with 100 µM NaAsO_2_ for 45 min. Scale bar = 20 µm. (B) Quantification of percent WT and DHX36hm U-2 OS cells containing 2 or more stress granules after treating with 100 µM NaAsO_2_ for t = 15, 30, 45 and 60 min. Data analyzed with two-tailed unpaired t-test and represented as mean ± s.d. (\**P* = 0.031, \*\**P* = 0.0011). Three biological replicates quantified. (C) Immunoblot of DHX36 and GAPDH in extracts from WT, DHX36hm and DHX36hm+WT cells, where +WT represents rescue with WT DHX36. (D-E) Quantification of percent WT, DHX36hm and DHX36hm+WT U-2 OS cells containing 2 or more stress granules after treating with 100 µM NaAsO_2_ for the times indicated. Data analyzed with one-way ANOVA, corrected with Tukey’s multiple comparisons test and represented as mean ± s.d. (\**P-adj* = 0.019 [WT vs DHX36hm, t = 45 min], \**P-adj* = 0.037 [WT vs DHX36hm, t = 60 min], \*\**P-adj* = 0.0098). Three biological replicates quantified. (F) Immunofluorescence of DAPI (blue) and PABPC1 (green) in WT and stable DHX36hm U-2 OS cells treated with 300 µM NaAsO_2_ for 30 min and recovered in DMEM for 45 min. Scale bar = 20 µm. (G) Quantification of WT and DHX36hm U-2 OS cells containing 2 or more stress granules after treating with 300 µM NaAsO_2_ for 30 min and allowed to recover in DMEM for t = 0, 30, 45 and 60 min. Data analyzed with two-tailed unpaired t-test, and represented as mean ± s.d.. Three biological replicates quantified. (H-I) Quantification of percent WT and DHX36hm U-2 OS cells containing 2 or more stress granules after recovering in DMEM for (H) t = 45 min and (I) t = 60 min. Data represented as mean of three biological replicates.

We further confirmed that this was a direct effect of DHX36 by overexpressing FLAG-DHX36 WT in DHX36hm cells (Fig. 4C). Overexpression of FLAG-DHX36 WT led to a reversal in the rate of stress granule formation, similar to that of WT levels (Fig. 4D & 4E).

We also examined if DHX36 affects stress granule disassembly after stress is removed. We treated cells with 300 µM NaAsO_2_ for one hour to ensure that stress granules were formed in all cells before removing the stress. Following arsenite removal, we observed that stress granules in DHX36hm cells persisted longer than in WT cells (Fig. 4F & 4G). Although not statistically significant, the trend of DHX36hm cells displaying a slower stress granule dissolution was consistent in our replicates at both 45 min and 60 min post-recovery (Fig. 4H & 4I).

Taken together, these two observations suggest that DHX36 can function to limit stress granule formation and/or promote disassembly.

### DHX36 reduction restores stress granule-like foci in G3BP1/2 KO cells

In an orthogonal approach to ascertain whether DHX36 can limit stress granule assembly, we knocked down DHX36 in cell lines lacking G3BP1 and G3BP2 (Supplementary Fig. 3A & 3B, middle and right). G3BP1 and its paralog G3BP2 are major assembly factors of stress granules. Stress granule formation is impaired in cells lacking these two proteins due to impaired G3BP1/2-mediated protein-protein and/or protein-RNA interactions (8). These cell lines therefore provide a sensitive assay to detect proteins that restrict stress granule formation via limiting RNA-RNA or RNA-protein interactions (31). If a protein such as the DEAD-box helicases eIF4A or DDX6 functions to limit stress granule formation through restricting RNA-mediated interactions, one can observe a restoration of small stress granule-like foci in G3BP1/2 dKO cells when eIF4A or DDX6 is functionally inhibited (31, 33).

Strikingly, we observed a rescue of stress granule-like foci in 31.7 ± 12.5% of G3BP1/2 dKO cells with a transient knockdown of DHX36 using siRNAs (Fig. 5A & 5B). In comparison, stress granules were observed in 8.5 ± 3.3% of cells with the non-targeting control, as visualised by PABPC1 staining (Fig. 5A & 5B) and oligod(T) RNA (Fig. 5A, right). Similarly, when we stably knocked down DHX36 in G3BP1/2 dKO cells with Cas9 targeting strategy, we also observed a rescue of stress granule-like foci in 18.1 ± 4.9% and 17.9 ± 1.4% of cells in two independent knockdown clones of DHX36 compared to that 8.7 ± 3.0% in G3BP1/2 dKO cells (Fig. 5C & 5D).

**Figure 5.**
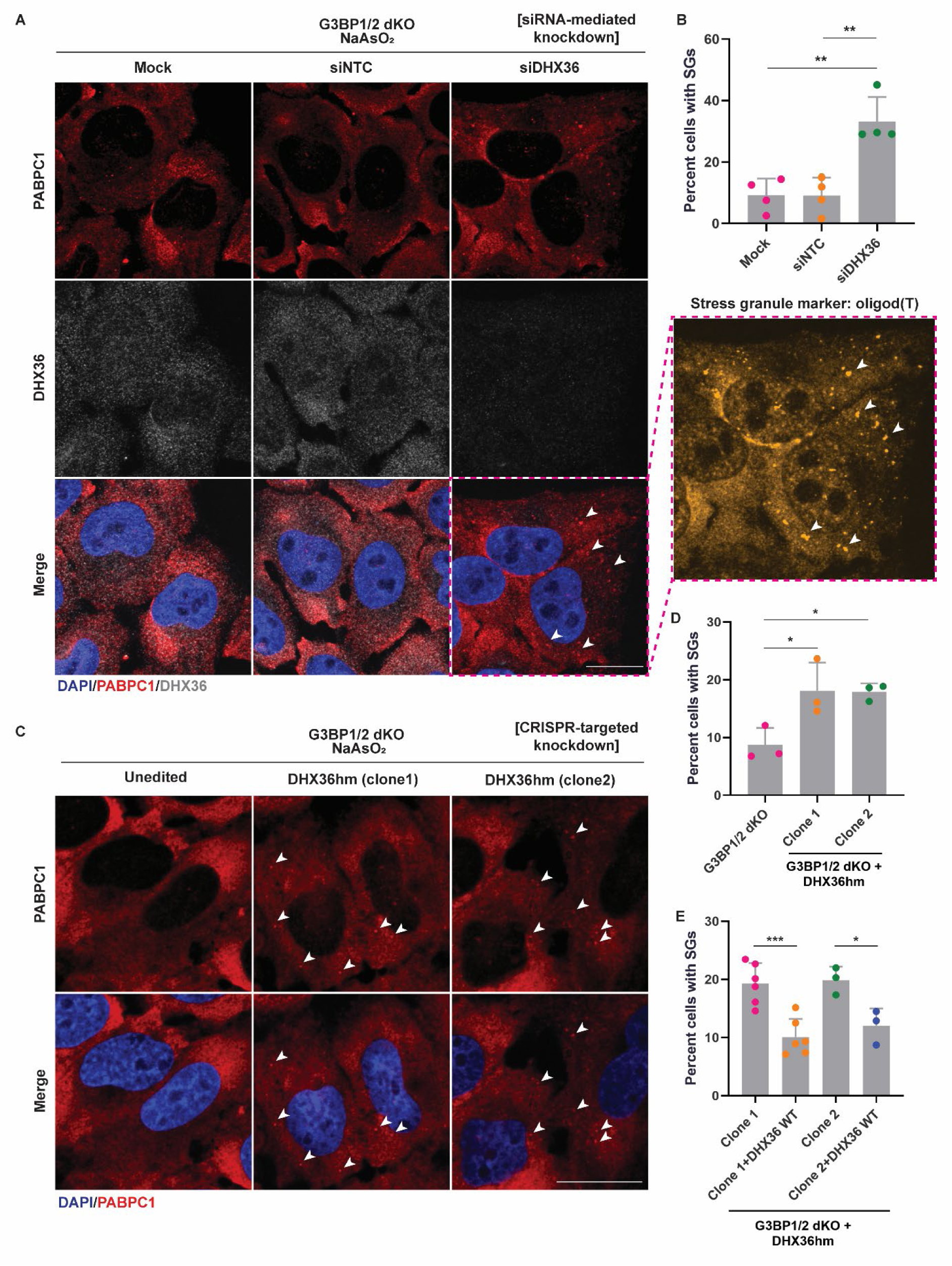
DHX36 reduction restores stress granule-like foci in G3BP1/2 KO cells. (A) Immunofluorescence of DAPI (blue), PABPC1 (red), oligo d(T) RNA (yellow) and DHX36 (grey) in G3BP1/2 dKO U-2 OS cells transfected with either siNTC (non-targeting control) or siDHX36. 48 hours post-transfection, cells were treated with 500 µM NaAsO_2_ for 60 min. White arrows: representative stress granule foci. Scale bar = 20 µm. (B) Quantification of cells with stress granules as shown in (A). Data analyzed with one-way ANOVA, corrected with Tukey’s multiple comparisons test, and represented as mean ± s.d. (\*\**P-adj* = 0.0014 [Mock vs siDHX36], \*\**P-adj* = 0.0014 [siNTC vs siDHX36]). Three biological replicates quantified. (C) Immunofluorescence of DAPI (blue) and PABPC1 (red) in G3BP1/2 dKO and G3BP1/d KO+DHX36hm U-2 OS cells treated with 500 µM NaAsO_2_ for 60 min. White arrows: representative stress granule foci. Scale bar = 20 µm. (D) Quantification of cells with stress granules as shown in (C). Data analyzed with one-way ANOVA, corrected with Tukey’s multiple comparisons test, and represented as mean ± s.d. (\**P-adj* = 0.033 [G3BP1/2 dKO vs Clone 1], \**P-adj* = 0.036 [G3BP1/2 dKO vs Clone 2]). Three biological replicates quantified. (E) Quantification of cells with stress granules in two independent clones of G3BP1/2 dKO+DHX36hm U-2 OS cells rescued with FLAG-DHX36 WT. Data analyzed with one-way ANOVA, corrected with Tukey’s multiple comparisons test, and represented as mean ± s.d. (\**P-adj* = 0.039, \*\**P-adj* = 0.0009). Six biological replicates quantified for Clone 1, three biological replicates quantified for Clone 2.

To validate that this restoration of stress granule-like foci was due to decreased DHX36 function, we performed a rescue experiment where we transduced FLAG-DHX36 WT into the G3BP1/2dKO+DHX36hm cell lines and confirmed DHX36 expression via immunoblot (Supplementary Fig. 5A). We showed that restoring DHX36 levels reversed the formation of stress granule-like foci in cells to 10.1 ± 3.1% and 12.0 ± 3.0% in two independent rescue cell lines (Fig. 5E, Supplementary Fig. 5B).

These observations suggest that DHX36 plays a role in limiting RNA condensation via disrupting RNA-RNA interactions either non-specifically or specifically on *trans* rG4s that form and can contribute to RNA condensation during stress.

## DISCUSSION

Herein, we present several observations arguing that rG4 motifs in *cis* do not affect mRNA partitioning into stress granules in U-2 OS cells. First, on the single mRNA level, we examined stress granule enrichment in both reporter and candidate endogenous mRNAs that contained rG4 motifs. The fraction of smFISH spots enriched in stress granules compared to that in the cytoplasm was independent of having rG4 sequences within the mRNA. Second, we found no correlation between having rG4 in an mRNA and its stress granule partitioning when we controlled for mRNA length. Third, when we further pushed the cellular equilibrium to favor rG4 formation by knocking down the DHX36 helicase or adding the rG4-stabilizing ligand cPDS, we observed no effect on stress granule partitioning of endogenous mRNAs with putative rG4s.

We consider two reasons why rG4 motifs alone might be insufficient to alter mRNA localization in stress granules.

First, the effect of rG4 motifs could be small and therefore be masked by other RNA features. This view is based on the fact that rG4 sequence motifs represent a small proportion of an mRNA and their effect, if any, could be masked by the rest of the mRNA. Compared to the length of a typical mRNA of 2 kb, each a rG4 motif (rG4 motif as defined by (G_≥2_X_1-7_G_≥2_X_1-7_G_≥2_X_1-7_G_≥2_, X is any other nucleotide) makes up only 0.8-2% of the entire mRNA. For a stress granule-enriched mRNA with an average length of 7.1 kb (5), this percentage is even lower. However, in our reporter mRNAs with five rG4 sequence motifs where the proportion of rG4 in the mRNA increased to around 15%, there remained no change in mRNA partitioning (Fig. 1C & 1E). While we do not rule out the possibility that rG4s can have as a small effect, we posit that the contribution of rG4 motifs to an mRNA’s stress granule partitioning is minimal compared with other features of the mRNA, such as hairpins and internal loops, interactions with RNA binding proteins and mRNA length.

A second issue is that the majority of rG4 structures is likely to exist in an unfolded state *in cellulo* and therefore would not impact stress granule partitioning. Guo & Bartel (40) failed to detect rG4 structures in human cells and suggested that the dynamic formation of rG4s is regulated by RNA binding proteins, particularly RNA helicases such as DHX36 (40, 41). In this study, we created a DHX36 hypomorph cell line with reduced DHX36 expression. We show that even when DHX36 levels are repressed, and RNA abundance is increased as shown previously (22), the abundance of endogenous rG4-containing mRNAs in stress granules remains unchanged. This could perhaps be due to a compensation by the RNA helicase repertoire in cells. While DHX36 is thought to be the predominant RNA helicase that unwinds rG4, other RNA helicases such as DDX1 (51), DDX21 (52) and DHX9 (53) have also been demonstrated to resolve rG4 structures. Thus, suppression of DHX36 could be compensated through functional redundancy with other RNA helicases. As such, we would not be able to observe a change in mRNA partitioning due to rG4s.

While DHX36 did not affect rG4-containing mRNA’s partitioning to stress granules, we did observe that DHX36 could affect stress granule formation. First, in our stress granule assembly and disassembly kinetics, we observe that cells depleted of DHX36 formed stress granules faster and dissolved stress granules more slowly than WT cells (Fig. 4). Second, in G3BP1/2 dKO cells that do not form canonical stress granules (8), a reduction in DHX36 expression, via either siRNA-mediated knockdown or Cas9 targeting, rescued the formation of stress granule-like foci in a subset of cells (Fig. 5). These observations indicate that DHX36 contributes to limiting formation of stress granules.

DHX36 might limit stress granules by two general mechanisms.

First, DHX36 could limit the formation of rG4s in *trans*, thereby reducing the number of intermolecular rG4s that stabilize the stress granule assembly. Alternatively, or in addition, DHX36 could limit other types of intermolecular RNA-RNA or protein-RNA interactions that promote stress granule formation.

In either case, the role of DHX36 in limiting stress granule formation extends support for the “RNA chaperone network” model where RNP granule assembly is thought to be regulated by a network of RNA binding proteins, especially RNA helicases, that work to influence various RNA-RNA and RNA-protein interactions (3). Previously, DEAD-box proteins eIF4A, DDX6 and the abundant monomeric RNA binding protein YB1 have been show to function as a type of RNA chaperone network to limit stress granule formation (26, 31, 33, 54). Our study suggests DHX36 also acts to limit stress granule formation, although future experiments will be required to determine the specific mechanisms of DHX36 function. In addition, it will be important to investigate if different human cell types have similar or distinct levels of DHX36 and other RNA helicases.

In summary, our study offers an alternative perspective on the functional role of rG4 sequence motifs in RNA granules and provides additional support for the RNA chaperone network in regulating stress granule formation. Considering the current perspectives on rG4’s role in RNA granules, additional rigorous studies will help evaluate the functional significance of rG4 in its native cellular contexts.

## DATA AVAILABILITY

The data underlying this article will be shared on reasonable request to the corresponding author.

## ACKNOWLEDGMENTS

We thank Dr. Nancy Kedersha and Dr. Paul Anderson for U-2 OS cell lines. We thank Theresa Nahreini (Cell Culture Facility, Department of Biochemistry, University of Colorado, Boulder, CO, USA; RRID:SCR_018988), Dr Annette Erbse (Shared Instruments Facility, Department of Biochemistry, University of Colorado, Boulder, CO, USA; RRID: SCR_018986) and Dr. Joseph Dragavon (BioFrontiers Institute Advanced Light Microscopy Core Facility, University of Colorado, Boulder, CO, USA; RRID: SCR_018302).

## AUTHOR CONTRIBUTIONS

Li Yi Cheng: Conceptualization, Formal analysis, Investigation, Methodology, Validation, Visualization, Writing-original draft, Writing-review & editing. Nina Ripin: Conceptualization, Formal analysis, Investigation, Methodology, Supervision, Writing-review & editing. Thomas R. Cech: Funding acquisition, Supervision, Writing-review & editing. Roy Parker: Funding acquisition, Supervision, Writing-review & editing.

