## Supplemental materials for "Impact of G-tract RNAs and the DHX36 helicase on stress granule composition and formation"

This PDF file includes:

Supplementary Figure 1-5

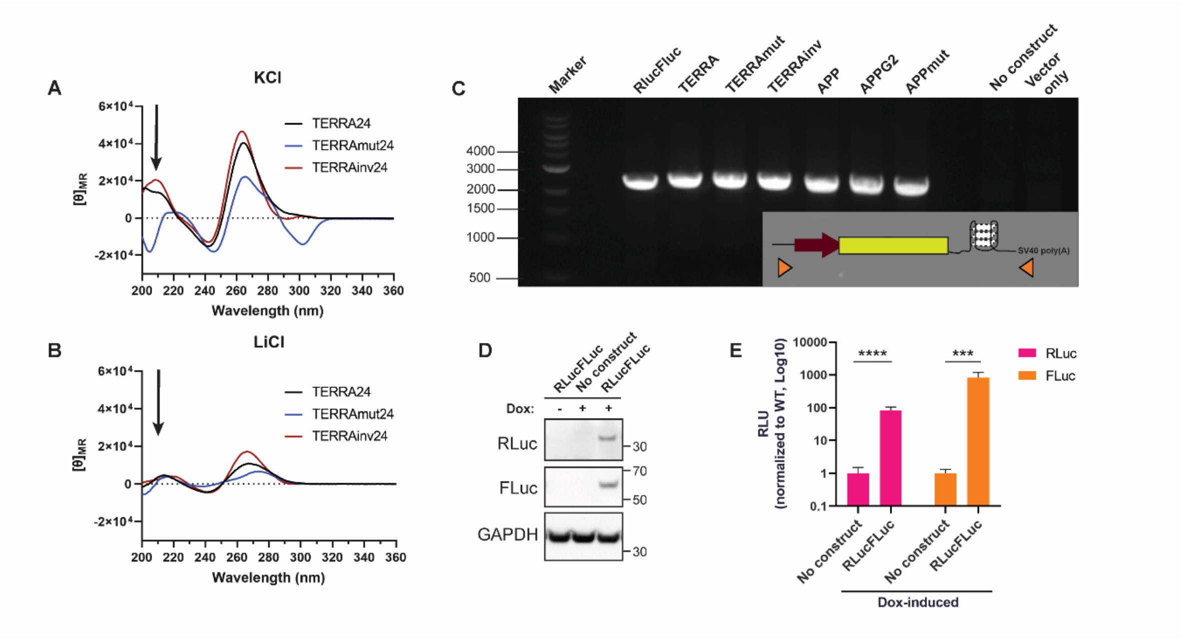


**Supplementary Figure 1** (A-B) Circular dichroism spectroscopy for TERRA24, TERRAmut24 and TERRAinv24 RNA in (A) 100mM KCl and (B) 100mM LiCl, at 25°C. Positive peak at 210 nm is indicative of RNA G4 structure while negative peak is indicative of A-form RNA, as shown by arrow. (C) DNA gel showing the integration of RLuc reporter into the AAVS1 locus using primers (orange triangle) flanking the reporter. (D) Immunoblot of RLuc, FLuc and GAPDH in extracts from WT and RLucFLuc U-2 OS cells induced with 4 μg/mL doxycycline. (E) Dual luciferase assay performed on WT and RLucFLuc U-2 OS cell extracts induced with doxycycline. Data analyzed with two-tailed unpaired t-test, and represented as mean ± s.d. (****P* = 0.0002, *****P* < 0.0001). Six biological replicates quantified.

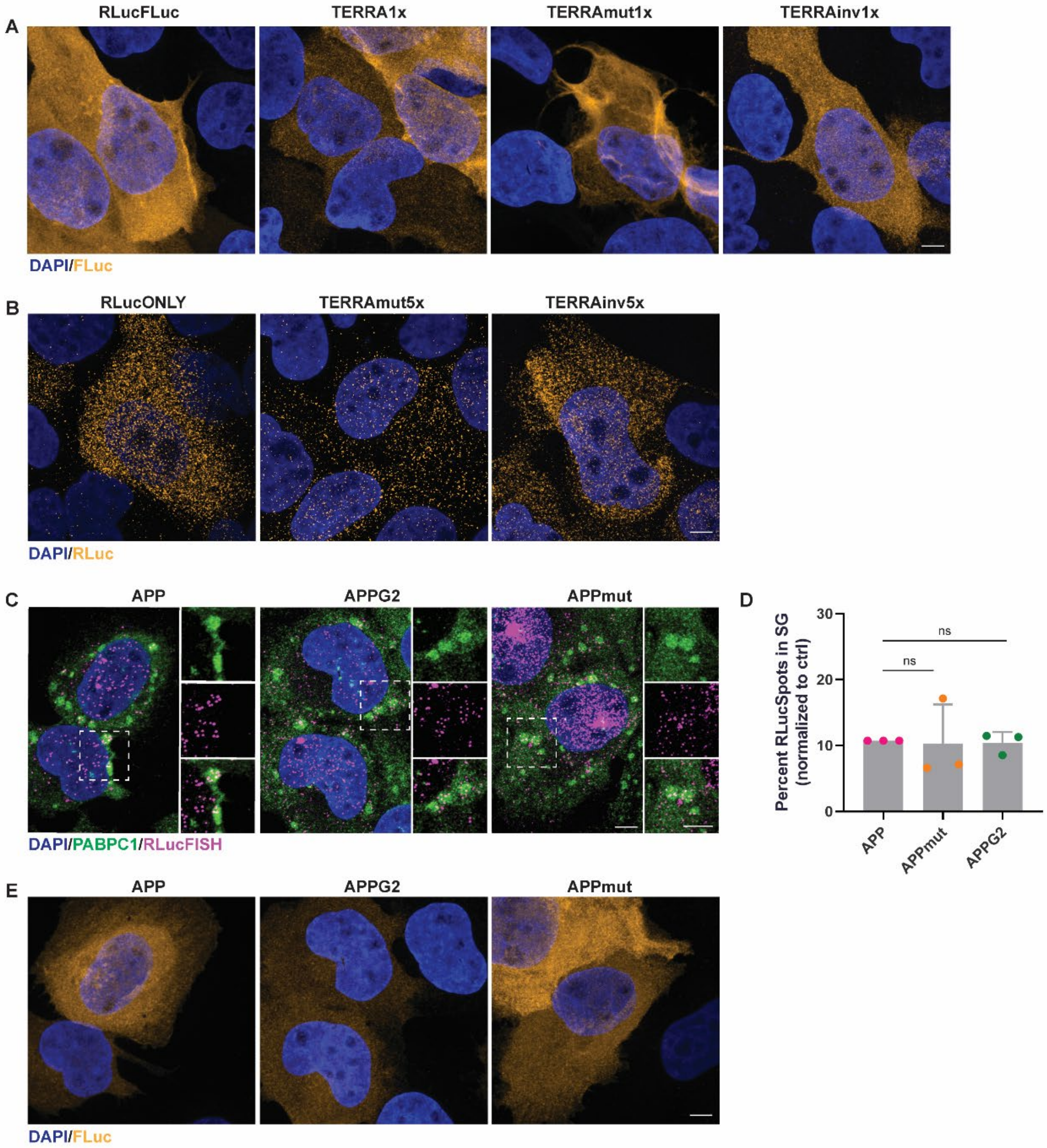


**Supplementary Figure 2** (A-B) Immunofluorescence of DAPI (blue) and luciferase (yellow) in U-2 OS cells stably expressing (A) 1x reporter constructs with FLuc and (B) 5x reporter constructs with RLuc. Cells are the same as those in Figure 1B and C. Scale bar = 5 µm. (C) Immunofluorescence of DAPI (blue), PABPC1 (green), and RLuc RNA (magenta) in U-2 OS cells stably expressing RLuc reporter constructs with APP sequences on the 3' UTR, previously described in (37). Scale bar = 5 µm. (D) Quantification of RLuc RNA FISH spots in stress granules as in (C), normalized to construct containing WT APP sequence. Data analyzed with one-way ANOVA, corrected with Tukey's multiple comparisons test and represented as mean ± s.d.. ns = non-significant, $p > 0.05$. Three biological replicates quantified. (E) Immunofluorescence of DAPI (blue) and FLuc (yellow) in U-2 OS cells stably expressing APP sequences. Cells are the same as those in (C). Scale bar = 5 µm.

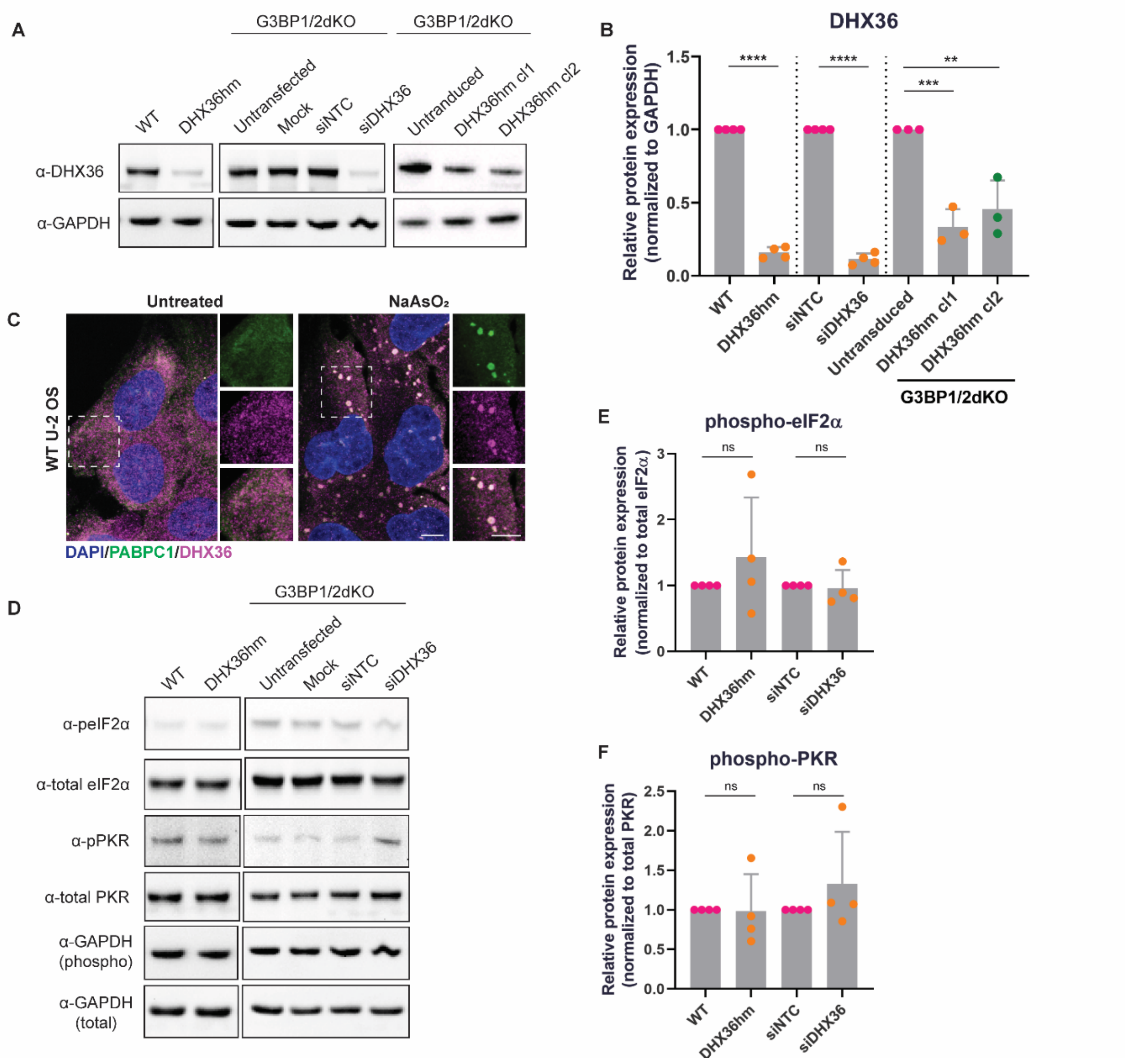


**Supplementary Figure 3** (A) Immunoblot of DHX36 and GAPDH in extracts from various U-2 OS cell lines. (B) Quantification of DHX36 expression in various cell lines normalized to GAPDH, and compared to their corresponding controls. For comparisons between two conditions, data analyzed with two-tailed unpaired t-test and represented as mean ± s.d. (*****P* < 0.0001). Four biological replicates quantified. For comparisons between three conditions, data analyzed with one-way ANOVA, corrected with Tukey's multiple comparisons test and represented as mean ± s.d. (***P-adj* = 0.0025, ****P-adj* = 0.0009). Three biological replicates quantified. (C) Immunofluorescence of DAPI (blue), PABPC1 (green) and DHX36 (magenta) in WT U-2 OS cells. Scale bar = 5 µm. (D) Immunoblot of phospho-eIF2α, total eIF2α, phosphor-PKR and total PKR in extracts from various U-2 OS cell lines. (E-F) Quantification of (E) phosphorylated eIF2α and (F) phosphorylated PKR expression in various cell lines normalized to total eIF2α and total PKR, respectively. Protein expression is compared to their corresponding controls. Data analyzed with two-tailed unpaired t-test and represented as mean ± s.d.. ns = non-significant, p > 0.05. Four biological replicates quantified.

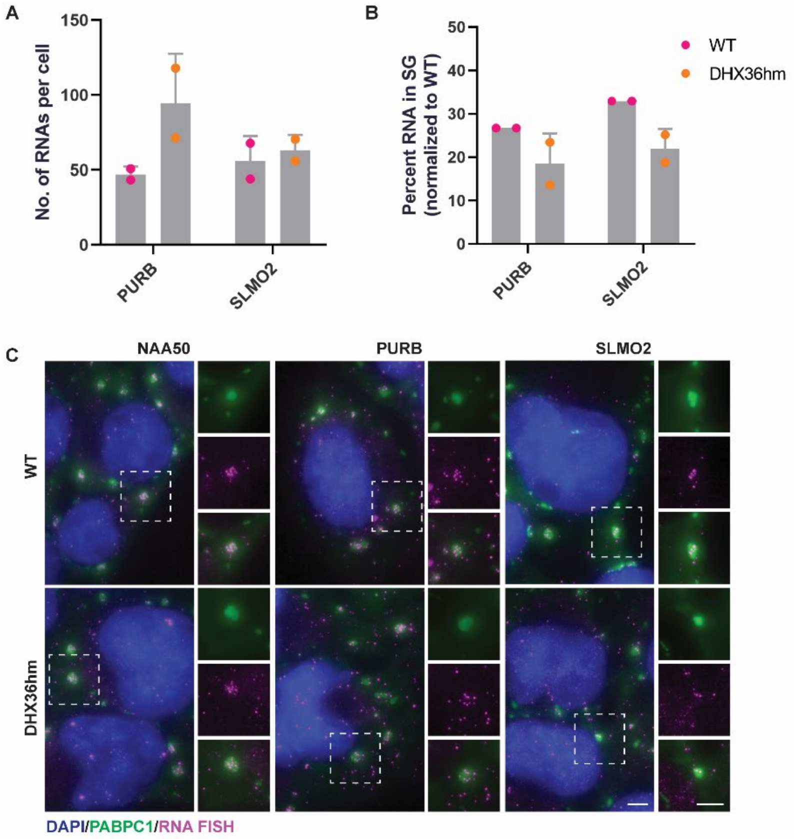


**Supplementary Figure 4** (A) Quantification of PURB and SLMO2 RNA FISH spots per cell. Data represented as mean ± s.d.. Two biological replicates quantified. (B) Quantification of percent PURB and SLMO2 RNA enrichment in stress granules. Data represented as mean ± s.d.. Two biological replicates quantified. (C) Immunofluorescence of DAPI (blue), PABPC1 (green) and NAA50, PURB and SLMO2 RNA (magenta) in WT and stable DHX36hm U-2 OS cells treated with 500 µM $NaAsO_2$ for 60 min. Scale bar = 5 µm.

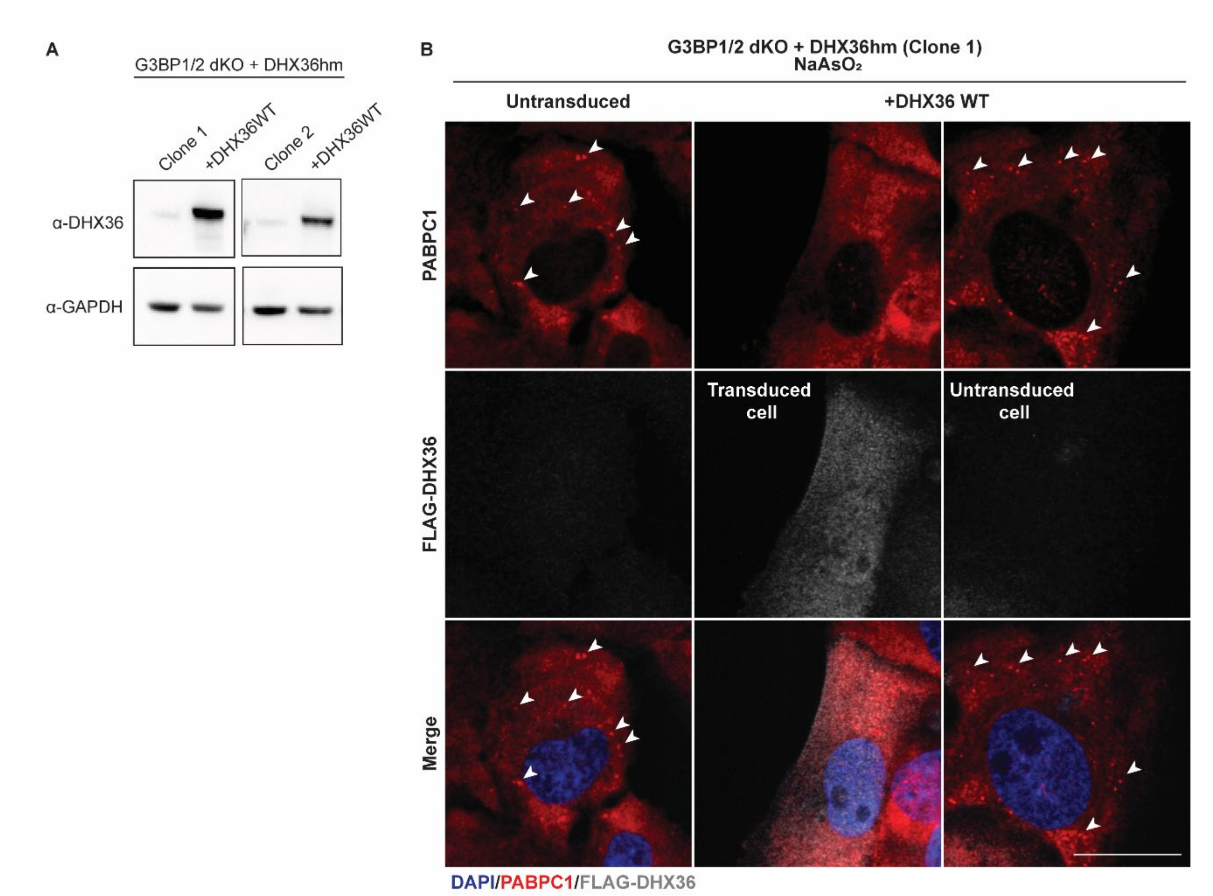


**Supplementary Figure 5** (A) Immunoblot of DHX36 and GAPDH in extracts from two independent clones of G3BP1/2 dKO+DHX36hm U-2 OS cells and rescued with FLAG-DHX36 WT. (B) Immunofluorescence of DAPI (blue), PABPC1 (red) and FLAG-DHX36 (grey) in G3BP1/2 dKO+DHX36hm U-2 OS cells and rescued with FLAG-DHX36 WT. Cells treated with 500 µM $NaAsO_2$ for 60 min. White arrows are indicative of stress granule-like foci. Scale bar = 20 µm.
